# Epigenetic Remodeling in Human Coronary Artery Smooth Muscle Cell Phenotypic Switching

**DOI:** 10.1101/2024.04.11.589030

**Authors:** Raja Chakraborty, Vincent P. Schulz, Kimberly Lezon-Geyda, Jui M. Dave, Kathleen A. Martin, Patrick G. Gallagher

**Author notes:** Co-Corresponding and co-senior authors; To whom correspondence should be addressed: Patrick G. Gallagher, AWRI, Nationwide Children’s Hospital, 575 Childrens Crossroads; Kathleen A. Martin, Yale Cardiovascular Research Center, 300 George St, Room 759, Columbus, OH 43054 New Haven, CT, 06511. These authors contributed equally to this work (Raja Chakraborty, Vincent P. Schulz).

## Abstract

**Background:** Smooth muscle cell (SMC) dedifferentiation contributes to repair and remodeling, but also cardiovascular pathologies. To understand this plasticity, the epigenetic landscape in SMC phenotypic switching was profiled.

**Methods:** Genome-wide analyses of histone modifications (ChIP-seq), chromatin architecture (ATAC-seq), and transcriptomes (RNA-seq) were performed on human coronary artery SMCs (CASMC) treated with rapamycin (contractile phenotype) and PDGF-BB (synthetic phenotype).

**Results:** Analyses of differentially acetylated promoter regions identified ZEB and ZBT7A as novel enriched regulatory motifs. There were more changes in the enhancer epigenome than in promoters in CASMC phenotypic switching. Rapamycin-activated enhancers were associated with differentiation and TGF-β signaling pathways and were most enriched in TEAD, SRF and SMAD motifs, whereas PDGF-induced enhancers were associated with ERK signaling and migration pathways, and were most enriched in ETV4, SOX5, and FOS motifs. GATA, TEAD, and SMCA1 motifs were enriched in CASMC enhancer open chromatin compared to other cell types. Candidate enhancers with single nucleotide polymorphisms linked to cardiovascular disease were markedly enriched in active enhancers and super enhancers and showed significant activity in reporter assays. In CASMC promoters and enhancers, common regulatory motifs were often enriched in both the differentiated and dedifferentiated phenotypes, suggesting that differential cofactor binding, as occurs with SRF at CArG elements, may be a more widespread mechanism underlying phenotypic switching.

**Conclusions:** These data identify novel regulatory elements engaged in SMC phenotypic switching and provide a comprehensive profile of SMC promoters, enhancers, super enhancers, and chromatin accessibility as a significant resource for studies of CASMC phenotype.

**Research Perspective:** *What Is New?:*

- This work identifies key regulatory elements and widespread changes in chromatin accessibility engaged in SMC phenotypic switching, including novel motifs enriched in promoters and enhancers. In CASMC promoters and enhancers, common regulatory motifs were often enriched in both the differentiated and dedifferentiated phenotypes, suggesting that differential cofactor binding, as occurs with SRF at CArG elements, may be a more widespread mechanism underlying phenotypic switching.
- This work identifies distinct enhancer profiles: enhancers activated by rapamycin were associated with TGF-β signaling and differentiation, while PDGF-induced enhancers were associated with ERK signaling and migration.
- Enhancer elements containing single nucleotide polymorphisms (SNPs) associated with cardiovascular disease from genome wide association studies (GWAS) showed notable enrichment across active enhancers and super enhancers. Select enhancers demonstrated statistically significant activity in reporter gene assays.

*What Question Should Be Addressed Next?:*

- The cis regulatory elements identified in this work suggest new transcription factors that can be tested to determine whether and how they may influence SMC phenotypic modulation.
- These studies could be extended to other stimuli to identify epigenomic signatures associated with CASMC transitions to other phenotypes including macrophages and chondrocytes.

## Introduction

In contrast to terminally differentiated skeletal and cardiac myocytes, mature smooth muscle cells (SMC) retain a unique plasticity that allows them to change their phenotype in response to environmental stimuli. This allows for the growth, repair, and remodeling of the vasculature, but also contributes to numerous vascular pathologies. In the context of vascular injury, such as occurs following angioplasty or surgical revascularization procedures, growth factors including PDGF-BB are released that induce SMC dedifferentiation from the contractile to a “synthetic” phenotype ^1,2^. During this dedifferentiation, SMC downregulate their SMC-specific repertoire of contractile proteins (which includes MYH11, TAGLN, CNN1, and ACTA2), and become proliferative, migratory, and secrete copious amounts of extracellular matrix (ECM) ^2,3^. This transition to a fibroblast-like phenotype allows for repair and remodeling of vessels after injury; however, this repair response often fails to resolve, leading to neointimal hyperplasia and restenosis^4^. Drug-eluting stents delivering rapamycin analogs have become a standard treatment for coronary artery angioplasty. Rapamycin is an mTOR Complex 1 (mTORC1) inhibitor. Although mTORC1 is a ubiquitously expressed and highly conserved master regulator of cellular anabolic processes ^1,5^, rapamycin has emerged as particularly effective in not only inhibiting SMC proliferation, migration, and protein synthesis, but in inducing the differentiated contractile program of gene expression ^1,6–9^.

Prior studies of SMC phenotypic switching have largely focused on proximal promoters of SMC-specific contractile genes, where CArG [CC(A/T)_6_GG] cis regulatory elements, which bind the transcription factor serum response factor (SRF) at SMC promoters, play a key role. The binding of distinct SRF accessory factors in response to environmental cues modulates SMC phenotypic plasticity. The co-activator Myocardin (MYOCD) binds to SRF at CArGs, enabling SMC-specific contractile gene expression and the differentiated state ^10^. In contrast, PDGF-BB induces KLF4 binding to G/C repressor elements near CArGs. KLF4 promotes dedifferentiation by inhibiting MYOCD binding at CArG motifs ^11^. TGFβ activation of Smad transcription factors also promote SMC differentiation, through binding Smad-binding elements (SBEs) and Smad interactions with myocardin and other transcription factors ^12,13^. Rapamycin has been shown to coordinately regulate myocardin-interacting factors, activating GATA6 ^7^ and inhibiting FoxO4 ^14^ to contribute to SMC differentiation.

The accessibility of DNA regulatory elements in promoters and enhancers determines their accessibility to transcription factors, with access to lineage-defining pioneer factors a key determinant of cell fate and identity. The epigenetic landscape is defined by functional changes to the genome not involving changes in the DNA sequence such as changes in chromatin architecture and DNA methylation. Dynamic alterations in chromatin architecture regulate numerous cellular processes with many chromatin functions regulated by histones ^15–18^. Histones are subject to numerous covalent modifications such as acetylation, phosphorylation, methylation, etc. that are central to the regulation of chromatin dynamics ^19–27^. These covalent modifications are closely linked to the regulation of transcription, thus the control of many biological processes, including CASMC plasticity ^28^.

Epigenetic regulation has been implicated in SMC plasticity, with reversible changes in DNA methylation and histone acetylation profoundly regulating gene expression. TET2, a methyl cytosine dioxygenase that opposes inhibitory DNA methylation, is a master epigenetic regulator of SMC phenotypic switching through regulation of key transcription factors, *MYOCD*, *SRF*, and *KLF4* ^1^. Conversely, DNA methyltransferases (DNMT) inhibit TET2 expression and promote SMC dedifferentiation, with DNMT inhibitors reducing intimal hyperplasia in mice ^29^. Histone acetylation is generally associated with gene activation, and histone deacetylases (HDACs) are induced by mitogens and associated with SMC intimal hyperproliferation ^3^. We have noted opposing roles for the histone acetyltransferases (HATs) p300 and CBP, with p300 promoting SMC differentiation, and CBP driving dedifferentiation ^6^. Notably, these epigenetic enzymes are regulated by stimuli that promote phenotypic switching. Rapamycin induces expression and activity of TET2 and p300, while PDGF inhibits their expression. Conversely, PDGF promotes CPB-dependent acetylation at genes associated with SMC dedifferentiation and recruitment of HDACs to contractile gene promoters ^6^.

Most studies of SMC phenotypic switching have focused on a small number of contractile gene promoters. However, enhancers contain the majority of binding sites for TFs that confer cell type specificity and dynamic patterns of gene expression ^30,31^. Therefore, a more thorough characterization of TF recognition motifs associated with cell type-specific enhancer repertoires could offer valuable insights into TFs that regulate SMC phenotypic switching. Studies have described the enhancers of endothelial cells (ECs), smooth muscle cells (SMCs), macrophages, and T cells, highlighting the dynamic changes in regulatory element activity when exposed to proatherogenic stimuli ^32,33^. Although these initial findings suggest enhancers have a substantial role in coordinating gene expression changes, a genome-wide comprehensive landscape of promoters, enhancers, and super enhancers using well defined promoter and enhancer marks has not been definitively established in SMC phenotypic modulation. We sought to bridge the gap in knowledge by profiling the epigenetic landscape in differentiated and dedifferentiated CASMCs.

## Methods

### Cell culture and treatment

Human coronary artery smooth muscle cells (hCASMCs, Promocell) were propagated in 199 medium (Gibco, 11150-067) supplemented with 10% FBS, 100 U/ml each penicillin-streptomycin, 2.7 ng/ml rhEGF (Biolegend, 713008), and 2ng/ml rhFGF (Biolegend 71034). Cells from passages 4 through 7 were used for all experiments. For rapamycin treatment SMCs were starved with 2.5% FBS for 24 hours. Following starvation, the cells were treated with Rapamycin (Sigma, R8781-200UL) with final concentration (50 nM), or with vehicle alone control (ethanol) for indicated time points. For PDGF-BB treatment, SMCs were treated with 0.5% FBS for 24 hours. Following starvation SMCs were treated with PDGF-BB (10ng/ml) (Sigma, P3201-50UG) and control untreated cells (PBS treated) for indicated time points. PDGF-BB is referred to as PDGF throughout the text. Genomic DNA (gDNA) and RNA were extracted from the same cells using the Qiagen DNA/RNA/miRNA all prep kit using manufacturer’s protocol after rapamycin or PDGF treatment with their individual vehicle ethanol and PBS treated control.

### RNA analyses

RNA was prepared for RNA-seq analyses as described ^34^. Samples were sequenced on an Illumina HiSeq 2500 using 75bp single-end reads. FASTQ format sequencing reads were aligned to the hg19 genome, NCBI Build 37, using STAR Version 2.5.3a software with default parameters and reads were counted in GENCODE v31 protein coding genes using Rsubread. The DESeq package was used to identify differences in expression with adjusted p-value <0.05. Gene Set Enrichment Analysis (GSEA) was performed using the clusterProfiler package ^35^.

### Chromatin immunoprecipitation (ChIP)

ChIP assays were performed as previously described ^36^. Samples were immunoprecipitated with antibodies against monomethyl histone 3 lysine 4 (H3k4me1, Abcam ab8895), trimethyl histone 3 lysine 27 (Abcam ab6002), acetyl histone 3 lysine 27 (H3K27Ac, Abcam ab4729), trimethyl histone 3 lysine 4 (Active Motif 39915), CTCF (Creative Diagnostics DMABT-H19813) and nonspecific rabbit IgG (sc-2091, Santa Cruz). ChIP was combined with the transposase based ChIPmentation protocol for library preparation ^37^.

### High throughput sequencing and data analyses

DNA processing and high throughput sequencing were performed as described ^34^. Sequenced reads were mapped to the human genome (hg19^38^ NCBI Build 37) using the BWA alignment program. The Model-based Alignment of ChIP-Seq (MACS2) program version 2.2.7.1 was used to identify narrow peaks in CTCF and H3K4me3 data. MACS2 was used to identify broad regions in H3K4me1 (--broad-cutoff 0.01) and H3K27Ac (--broad-cutoff 0.01 with fold change >4) ^39^. SICER version 1.1 was used to identify broad regions bound by H3K27me3 with FDR cutoff 0.01. Changes in histone modifications or open chromatin in different sample types were identified with the DiffBind package using full library size normalization. Localization of histone modifications relative to known genes was done using the ChIPseeker package. Motif finding was performed using the HOMER algorithm (Hypergeometric Optimization of Motif EnRichment) ^30^. The Genomic Regions Enrichment Annotations Tool (GREAT) was used to analyze functional significance of *cis*-regulatory regions identified by ChIP-seq ^40^. Smooth muscle cell specific regions of open chromatin were identified by merging peaks of different cell types, splitting into bins of 500 bases, and counting reads in each bin for each sample. Regions with 10-fold increased open chromatin with FDR 0.1 were identified using DESeq2. Super enhancers were identified using the ROSE algorithm.

### Identification and analysis of biologically relevant SNPs

The locations of SNPs shown to demonstrate highly significant linkage to cardiovascular-related traits were obtained from the catalog of published human Genome-Wide Association Studies (GWAS) compiled by the National Human Genome Research Institute and the European Bioinformatics Institute^41^. Using BedTools software, enhancers were intersected with cardiovascular-related trait SNPs and overlap identified.

### Reporter gene assays

Candidate enhancer regions near the *JPH2*, *CDKN2B*, *ELL*, *SLC1A1* and *PDGFD* genes were amplified using primers flanking the boundaries of called peaks (Supplemental Table S1) and cloned upstream of an SV40 promoter-firefly luciferase reporter cassette in the pGL3Promoter plasmid. Transfections into CASMC were performed as described ^7^ using 6ug pGL3 firefly luciferase reporter and 2ug *Renilla* luciferase transfection control plasmid). A dual Luciferase assay (Promega, #E2920) was used to measure the luciferase activity in transfected cells followed by normalization for each sample.

### Data access

The raw data files generated by the RNA seq, ATAC-seq, and ChIP-seq analyses have been submitted to Gene Expression Omnibus (GEO, http://www.ncbi.nlm.nih.gov/geo/, reference series GSE261841.

## Results

### Transcriptome analyses of human coronary artery smooth muscle cells (hCASMC) treated with rapamycin or PDGF by RNA-seq

hCASMCs were treated with rapamycin (50 nM) or PDGF-BB (10 ng/ml) for 48 hours as described ^6^ to compare gene expression and the epigenetic landscape in the distinct differentiated contractile and dedifferentiated synthetic phenotypic states, respectively. RNA was isolated, libraries prepared, and RNA-seq performed to obtain the transcriptomes (Supplemental Table S2). Multidimensional scaling, performed on expressed genes to assess sample relatedness, revealed rapamycin-and PDGF treated cells clustered apart, indicating that treatment led to distinct transcriptome composition. Differentially expressed transcripts among rapamycin and PDGF treated cells were identified, sorted by *p*-value in their expression (Supplemental Table S3) and differentially expressed genes displayed in heat maps (Figure 1A). Gene ontology analyses reveal varying patterns of gene expression that align with known functions of the differentiated and dedifferentiated SMC states. In rapamycin treated cells, enriched categories included translation and metabolism-related terms (Figure 1B), consistent with rapamycin’s potent and selective inhibition of mTORC1 and its well-known role as a central regulator of anabolic processes ^5^. Genes related to migration and chemotaxis were among the highest ranked categories for PDGF (Figure 1B) consistent with PDGF’s regulation of the reparative synthetic phenotype. These data indicate differing programs of gene expression in CASMCs after treatment to alter differentiation status. Gene Set Enrichment Analysis (GSEA) of gene expression in rapamycin and PDGF-treated smooth muscle cells yielded similar results (Figure 1C and 1D). Representative examples of genome browser tracks for genes differentially expressed with phenotypic switching include the contractile gene transgelin, *TAGLN*, induced after rapamycin treatment, and the membrane receptor beta integrin 1, *ITGB1,* induced after PDGF treatment^42^ (Figure 1E). These transcriptomic changes validate the cell culture model in which to evaluate corresponding changes in epigenetic regulation.

**Figure 1.**
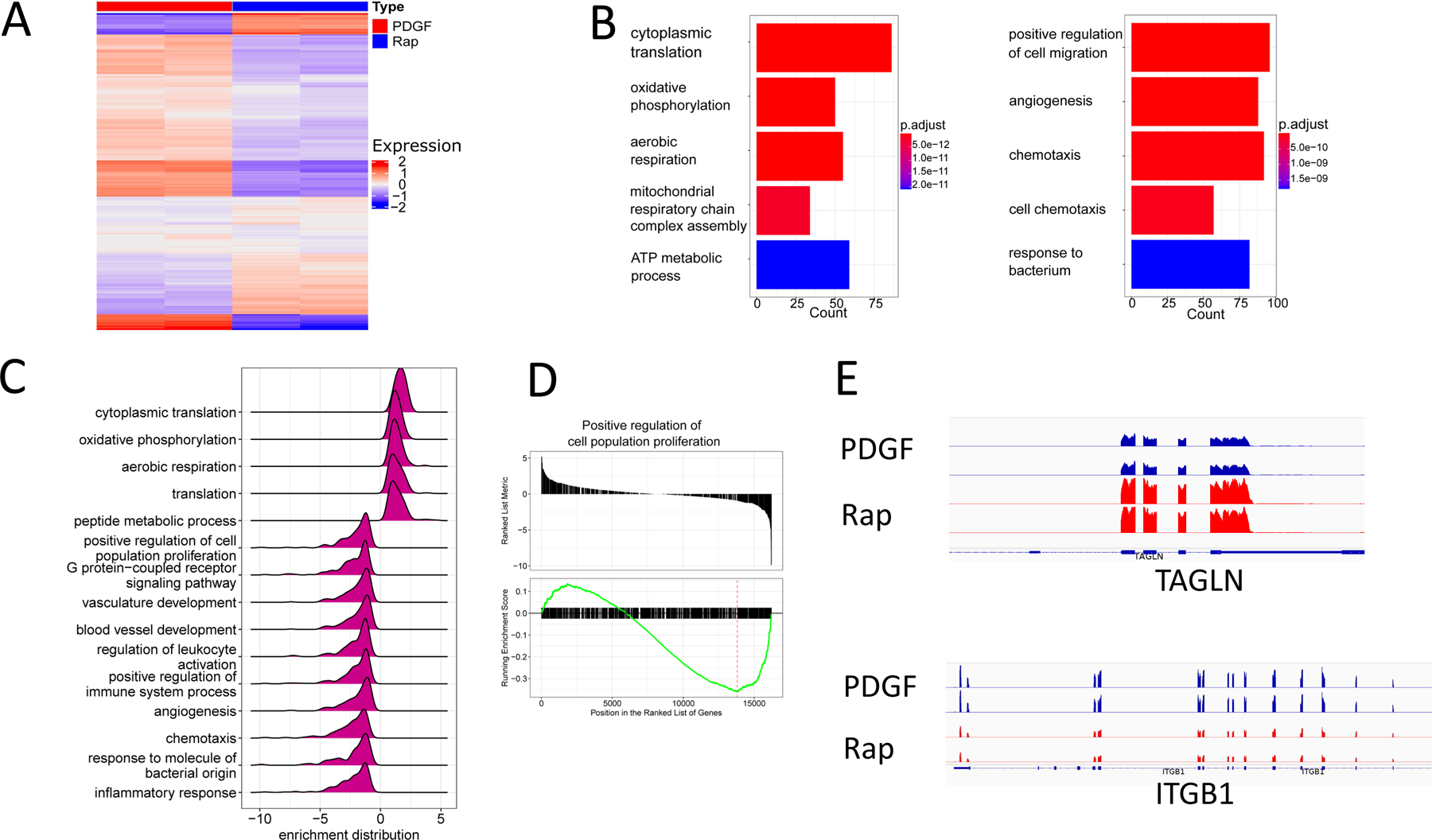
Transcriptome analyses: Human CASMCs treated with rapamycin or PDGF have distinct gene expression profiles. **A**. Heat map display of gene expression patterns of differentially expressed genes comparing rapamycin or PDGF-treated cells. Red represents elevated expression while blue represents decreased expression, compared with the row mean. Each column represents a biologic replicate. Genes displayed were selected based on fold changes of 2 or more and FDR adjusted *p* value < 0.05 between cell types. **B**. Categories of differentially expressed genes increased in rapamycin-treated cells. **C**. Categories of differentially expressed genes increased in PDGF-treated cells. **D.** Gene set enrichment analysis comparing rapamycin-versus PDGF-treated cells. **E**. Running enrichment score analysis of the angiogenesis gene set comparing rapamycin-versus PDGF-treated cells. **F.** RNA coverage profiles of representative differentially expressed genes.

### Identification of promoters differentially regulated by rapamycin and PDGF in CASMCs

We identified GENCODE predicted promoters marked by H3K4me3 and classified them as active (H3K4me3+, H3K27me3-) or poised (H3K4me3+, H3K27me3+). A heat map of promoters by class after treatment with rapamycin or PDGF is shown in Figure 2A. Changes in promoter state after rapamycin or PDGF treatment were examined (Figure 2B). Most promoter classes were conserved between rapamycin and PDGF, but a substantial number of promoters changed after treatment. An interesting finding was the presence of changes between poised and active promoter states in SMC phenotypic switching, as poised promoters, which are marked simultaneously by both active and repressive marks, are typically observed in stem cells and during development, cancer initiation and response to therapy, and are not typically described in differentiated somatic cells ^43^. Poised promoters are inactive despite the presence of both H3K4me3 and H3K27me3 marks ^44^.

**Figure 2.**
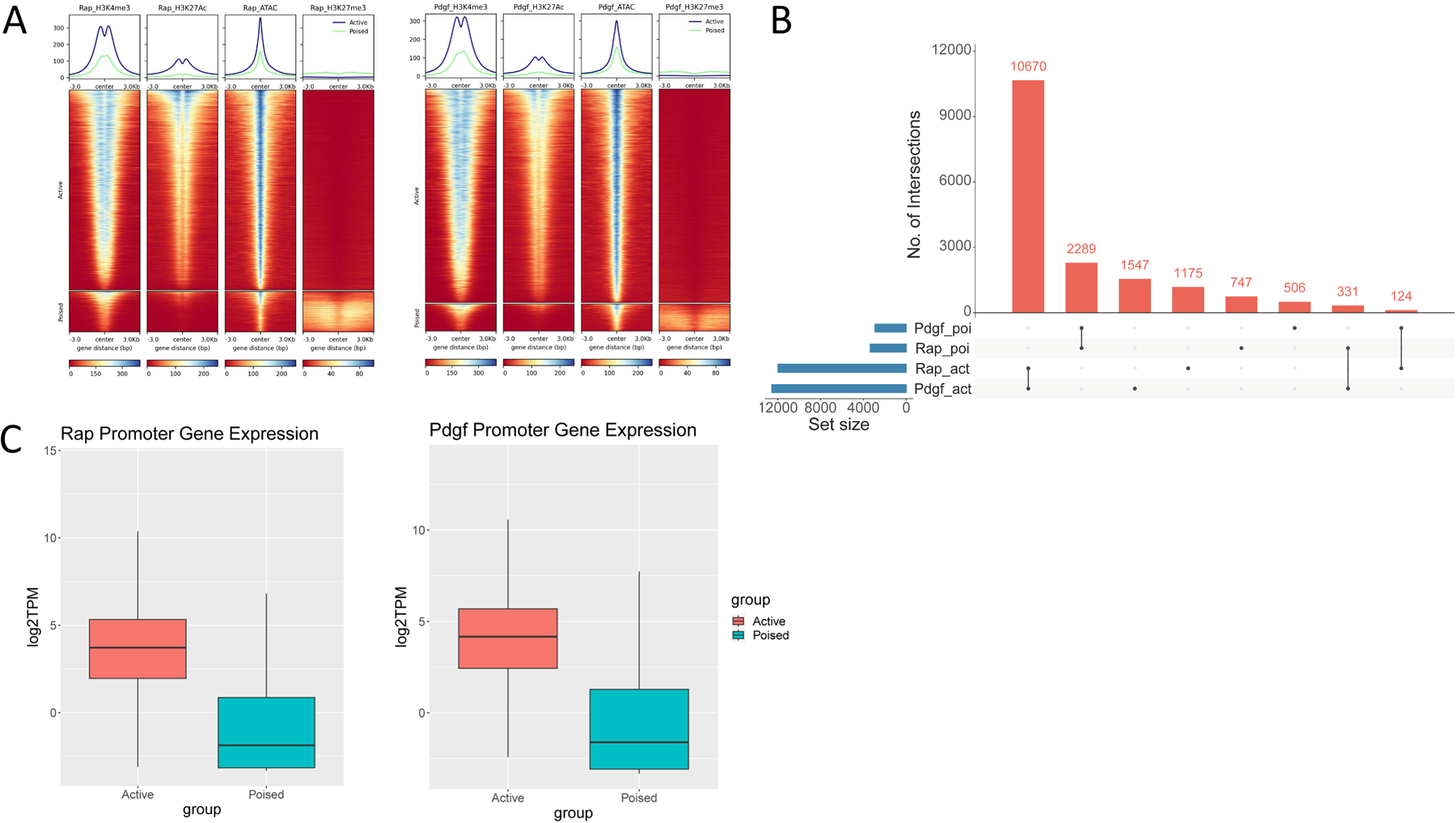
Promoter classes in rapamycin- and PDGF-treated CASMCs. A. Heat map of promoter classes active and poised after rapamycin or PDGF treatment. Signals of H3K4me3, H3K27Ac, ATAC-seq, and H3K27me3 centered over promoters are shown. B. Upset plot showing overlap in regions from rapamycin and PDGF promoter classes. Most promoter classes are conserved between rapamycin and PDGF, but 331 promoters change from PDGF active to rapamycin poised state and 124 promoter change from rapamycin active to PDGF poised state. C. Boxplots of gene expression and promoter class in rapamycin- and PDGF-treated cells.

When comparing rapamycin-to PDGF-treated cells, the largest number of changes were from PDGF active promoter state to rapamycin poised promoter state (331 promoters), followed by rapamycin active promoter state to PDGF poised promoter state (124). Similarly, when comparing control cells to rapamycin treated cells, 146 promoters changed state from active to poised promoters and 194 promoters changed state from poised to active after rapamycin treatment. Linked gene expression was much higher in active promoters than poised promoters (Figure 2C). Despite these associations, promoters changing between poised and active states represented a minor fraction of differentially expressed genes, and pathway analysis of promoters in this category did not reveal genes typically associated with the contractile or synthetic phenotypes.

Active promoters and their linked gene expression were analyzed to assess regulation of genes that are highly expressed in CASMCs compared to other cell types. The top 500 CASMC expressed genes were identified after removing housekeeping and ubiquitously expressed genes (200 genes with greatest expression across multiple cell types in the GTEx RNA-seq dataset ^45^) in the rapamycin- and PDGF-treated states (Supplemental Table S4). These included genes typically associated with phenotypic switching, such as *ACTA2*, *TAGLN*, *TGFBI, and MYL12A* (differentiated) and *MMP1, SERPINE1, CXCL12, THBS1,* and *TIMP1* (dedifferentiated). To identify key regulatory promoter elements associated with gene expression, the HOMER algorithm ^30^ was utilized to identify transcription factor motifs centered on the active promoter-associated peak of open chromatin. Enriched motifs included well known regulators of the SMC phenotype, such as KLF and SRF for both differentiated and dedifferentiated cells. There was considerable overlap between the enriched promoter motifs found in the differentiated and dedifferentiated SMC states, suggesting combinations of transcription factors are responsive to extracellular signals to rapidly change promoter activity at hallmark genes in phenotypic switching. A classic example is the SRF-binding CArG motif that is recognized as a key regulatory element driving contractile genes that can be inhibited with growth factor stimulation by recruiting distinct ternary complex factors that replace myocardin^46^. Our data suggest multi-functional regulatory motifs may be a common characteristic of SMC promoters that undergo phenotypic switching.

In addition to marking active enhancers, H3K27Ac is often found at sites of active promoters, and levels of acetylation are associated with levels of linked gene transcription^47^. Promoters marked by differential patterns of H3K27 acetylation in CASMCs after treatment with rapamycin or PDGF were identified and displayed in heat maps (Figure 3A). To analyze regulatory elements, we identified motifs enriched in differentially acetylated regions centered on peaks of open chromatin. Enriched motifs identified with differential H3K27Ac in rapamycin-treated cells compared to PDGF treated cells included ZEB, ZBT7A, and E2F5 (Figure 3B). ZBT7A and ZEB were also enriched in PDGF-treated cells compared to rapamycin treated cells, as well as GATA (Figure 3B). Categories of biological processes enriched in promoters with differential patterns of H3K27Ac matched known functions of the differentiated or dedifferentiated phenotypes, with rapamycin-associated pathways including cytoskeleton and muscle contraction whereas promoters with differential patterns of H3K27Ac after PDGF treatment included chemotaxis and migration (Figure 3C). We then investigated whether the distribution of H3K27Ac around the transcription start site (TSS) correlated with gene expression levels. We consistently observed that peaks of H3K27Ac overlapped with gene transcription start sites and patterns of H3K27 acetylation strongly correlated with levels of gene expression in both rapamycin- and PDGF-treated CASMCs. This relationship is illustrated in the genome browser tracks showing differential patterns of H3K27 acetylation at two well established SMC phenotypic markers: *CNN1* promoter, activated after rapamycin treatment, and the *S1PR1* (a gene that induces intimal hyperplasia ^48^promoter, activated after PDGF treatment (Figure 3D). These findings highlight a strong correlation between differential promoter H3K27 acetylation and gene activity with SMC phenotypic plasticity.

**Figure 3.**
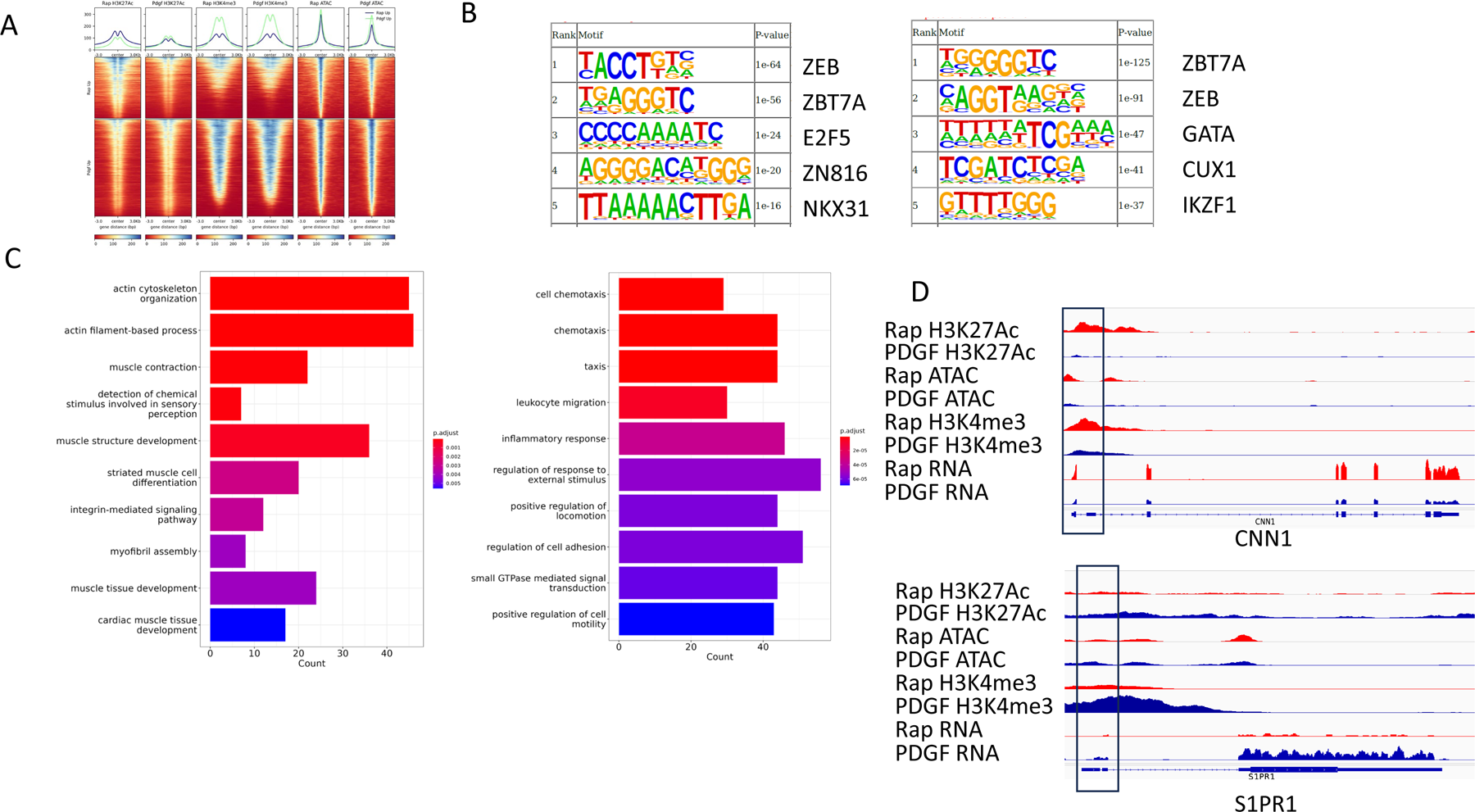
Promoters marked by differential patterns of histone modifications and regions of open chromatin in CASMCs after treatment with rapamycin or PDGF. A. Heat map of differential promoters after rapamycin or PDGF treatment. Regions increased in rapamycin treated cells are shown on top. PDGF treated cells are shown on the bottom. The signal density of H3K27Ac, H3K4me3, and open chromatin determined by ATAC-seq is plotted centered on the ATAC peak. B. Enriched motifs identified with differential H3K27Ac centered on the ATAC peak in rapamycin-treated cells. Right. PDGF-treated cells. C. Overrepresented GO biological processes. Left: rapamycin-treated cells; right: PDGF-treated cells. D. Genome browser tracks of differential promoters. Top. The *CNN1* gene promoter activated in rapamycin treated cells. The *S1PR1* gene promoter activated in PDGF-treated cells.

### Identification of multiple classes of enhancers in rapamycin and PDGF-treated CASMCs

The influence of rapamycin and PDGF on the enhancer landscape in SMCs was examined. Enhancers are typically marked by histone 3 lysine 4 monomethylation (H3K4me1) and further classified as active (H3K4me1+, H3K27Ac+, H3K27me3-), intermediate (H3K4me1+, H3K27Ac-, H3K27me3-) or poised (H3K4me1+, H3K27Ac+/-, H3K27me3+), based on chromatin architecture, conservation, genomic location, levels of gene expression of associated genes, and predicted function of associated genes ^49,50^. Active enhancers are dynamic regulators controlling levels of expression of linked genes. Poised enhancers often regulate gene expression in response to various cellular stimuli, *e.g.*, differentiation cues, and are generally associated with genes with specialized functions. Intermediate enhancers are associated with genes involved in a wide variety of biological processes unlinked to cell type. To identify enhancer classes in rapamycin and PDGF-treated SMCs, genome-wide maps of histone architecture were created using ChIP-seq with antibodies for H3K4me1, H3K27me3, and H3K27Ac using CASMC chromatin. To exclude gene promoters, regions with an H3K4me1 peak located within 1kb of annotated transcriptional start sites (TSS) and an H3K4me3 peak were excluded from the analyses. There were 32246 active, 52151 intermediate, and 7107 poised enhancers in rapamycin-treated cells and 35732 active, 55549 intermediate, and 4502 poised enhancers in PDGF-treated cells (Supplemental Table S5). Heat map analyses revealed patterns of distinct chromatin architecture characteristic of each of the enhancer classes (Figure 4A). Active enhancers demonstrated high levels of H3K27Ac and open chromatin with no repressive marks. Intermediate enhancers demonstrated low levels of H3K27Ac, no repressive marks, and relatively low levels of open chromatin. Poised enhancers demonstrated low levels of H3K27Ac, high levels of H3K27me3, and medium levels of open chromatin (Figure 4A). The human genome was portioned into seven bins relative to GENCODE genes corresponding to promoter, 5’ UTR, exons, introns, downstream, and intergenic regions. Sites of specific histone modifications and enhancer class were assigned to these bins and percentages calculated (Figure 4B). In both rapamycin and PDGF-treated chromatin, active and intermediate enhancers were mainly in intronic regions, whereas poised enhancers were found more frequently in intergenic regions. Comparing enhancer states between rapamycin- and PDGF-treated cells, many enhancers had the same state in both rapamycin and PDGF-treated SMCs. Notably, some enhancers changed state from intermediate to active after rapamycin or PDGF treatment (Figure 4C, D), along with changes in key biological processes shown in (Figure 4E), which could reflect a mechanism for rapid SMC phenotypic switching in response to extracellular cues.

**Figure 4.**
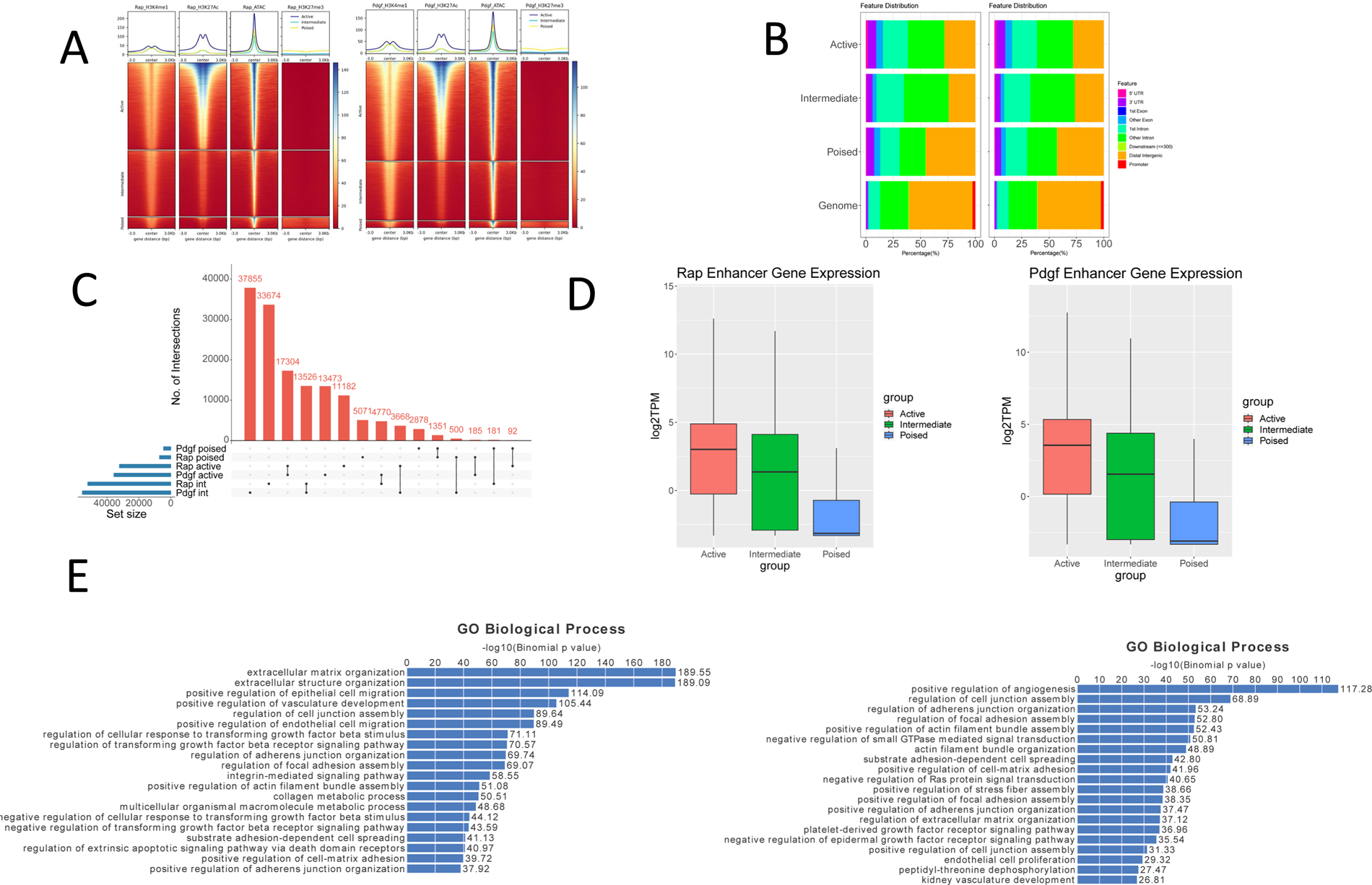
Histone modification density and enhancer class in CASMCs chromatin after treatment with rapamycin or PDGF. The signal density of H3K4me1, H3K27Ac, and H3K27me3 chromatin and ATAC-seq signal is plotted relative to the ATAC peak. **A.** Rapamycin-treated smooth muscle cells. B. PDGF-treated smooth muscle cells. C. Location of enhancers relative to annotated gene features, e.g. promoter, intron, etc. D. Upset plot of active and intermediate enhancers demonstrating some enhancers have the same state in both rapamycin and PDGF-treated cells and some enhancers change state from intermediate to active after rapamycin or PDGF treatment. E. Boxplots of nearest gene expression and associated enhancer class in rapamycin and PDGF-treated cells.

### Expression and function of genes associated with enhancer class in rapamycin and PDGF-treated CASMCs

Cell and tissue-type specific enhancers typically act over distances of tens to hundreds of kilobases ^51^. Thus, *bona fide* CASMC enhancers are expected to be enriched in the vicinity of genes that are expressed and functional in their respective cells ^52–54^. To determine whether CASMC enhancers are localized in this manner, gene expression in rapamycin and PDGF treated CASMCs was correlated with each enhancer class. Levels of gene expression of associated genes were highest for active enhancers and lowest for poised enhancers, with levels of expression between these values for intermediate enhancers in both rapamycin and PDGF-treated CASMCs, suggesting that enhancer state is a strong determinant of SMC gene expression.

### Differential patterns of enhancer H3K27 acetylation in CASMCs after treatment with rapamycin or PDGF

To identify enhancers differentially regulated in CASMC phenotypic states, enhancers marked by differential patterns of H3K27 acetylation after treatment with rapamycin or PDGF were identified and heat maps prepared with the signal density of H3K27Ac, H3K4me1, and ATAC-seq plotted centered on the ATAC peak (Figure 5A). Similar patterns of increased H3K27Ac and open chromatin were observed in differential regions from rapamycin- and PDGF-treated CASMCs, in parallel with little to no changes in H3K4me1. Motif analysis of regions with increased acetylation identified the most highly enriched transcription factor motifs in enhancers. In rapamycin treated cells, the top motifs included TEAD4, SRF, and SMAD4 whereas the motifs most enriched in PDGF-treated cells included ETV4, SOX5, and FOS (Figure 5B). We performed a statistical enrichment analysis of GO biological process terms associated with enhancers exhibiting differential patterns of H3K27 acetylation (Figure 5C). For enhancers activated by rapamycin treatment, top enriched terms were terms associated with TGFβ signaling and differentiation. In contrast, enhancers induced by PDGF treatment showed enrichment for terms linked to growth factor, MAPK and inflammatory signaling. There was considerable overlap in pathways regulated by rapamycin and PDGF, although many of these were regulated in opposite directions, as expected, in the contractile versus synthetic states. These included response to wound healing, extracellular matrix and actin cytoskeleton organization, proliferation, and migration. An enhancer near the *SMAD2* gene is an example of an enhancer with differential patterns of H3K27Ac with phenotypic switching. The larger H3K27Ac peak at this enhancer with rapamycin treatment is associated with increased RNA transcripts relative to the PDGF-treated condition (Figure 5D). Identification of differentially regulated enhancer motifs suggests several transcription factors and signaling pathways for further investigation as these may be important mediators of phenotypic switching.

**Figure 5.**
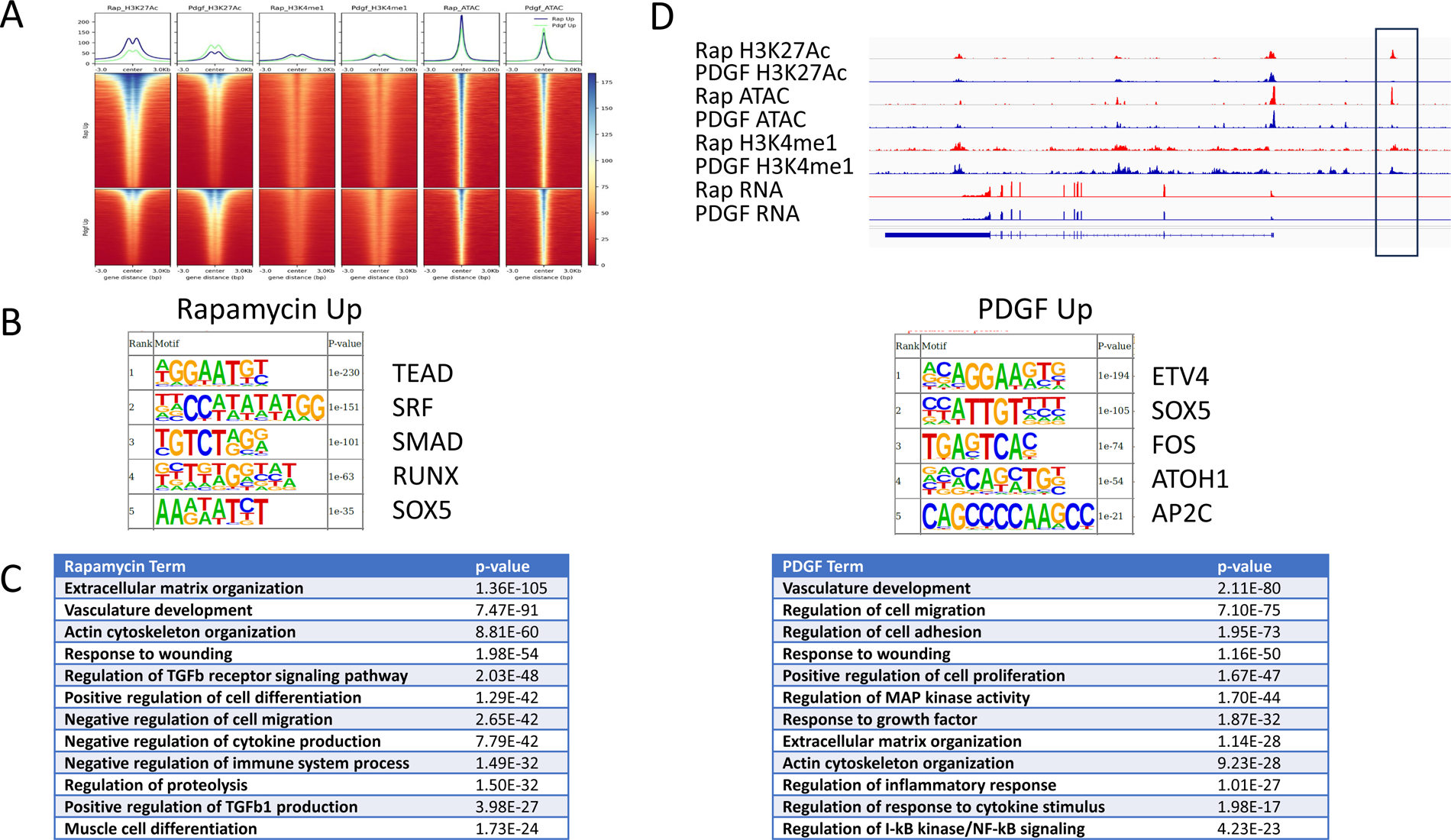
Enhancers marked by differential patterns of histone 3 lysine 27 acetylation in CASMCs after treatment with rapamycin or PDGF. A. Heat map of differential enhancers after rapamycin or PDGF treatment. Regions increased in rapamycin treated cells are shown on top. PDGF treated cells are shown on the bottom. The signal density of H3K27Ac, H3K4me1, and ATAC-seq is plotted centered on the ATAC peak. B. Motifs enriched in regions with increased acetylation in rapamycin treated cells. C. Enriched GO biological process terms associated with increased acetylation in rapamycin and PDGF treated cells. D. Genome browser track of a differential enhancer near the *SMAD2* gene in rapamycin-treated cells.

### Human coronary artery smooth muscle cells exhibit a distinct pattern of chromatin accessibility

One goal of our studies was to identify regions of open chromatin unique to CASMCs as markers of critical cell type-specific regulatory elements. To do this, we examined patterns of non-promoter chromatin accessibility in CASMCs and compared them to patterns in other muscle and non-muscle cell types (Figure 6A). CASMCs demonstrated distinct patterns of chromatin accessibility compared to other tissues, including other types of smooth muscle (visceral and reproductive tissues), other muscle types (cardiac and skeletal), and other vascular cell types such as fibroblasts, keratinocytes, and endothelial cells. DESeq2 read count analysis identified 3387 non-promoter ATAC peaks unique to CASMCs, displayed as a heat map in Figure 6B. Functional analyses using the Genomic Regions Enrichment of Annotations Tool (GREAT) algorithm of non-promoter differential ATAC peaks revealed enrichment for the mouse phenotype term “abnormal vascular smooth muscle morphology”. Motif analysis identified motifs enriched in regions of CASMC-specific open chromatin including GATA, TEAD, and SMCA1 (Figure 6C). An example of a CASMC-specific intronic enhancer ATAC peak at the *SMAD5* gene locus is shown (Figure 6D). These data suggest cis-regulatory elements in unique open chromatin regions are likely determinants of CASMC-specific gene expression.

**Figure 6.**
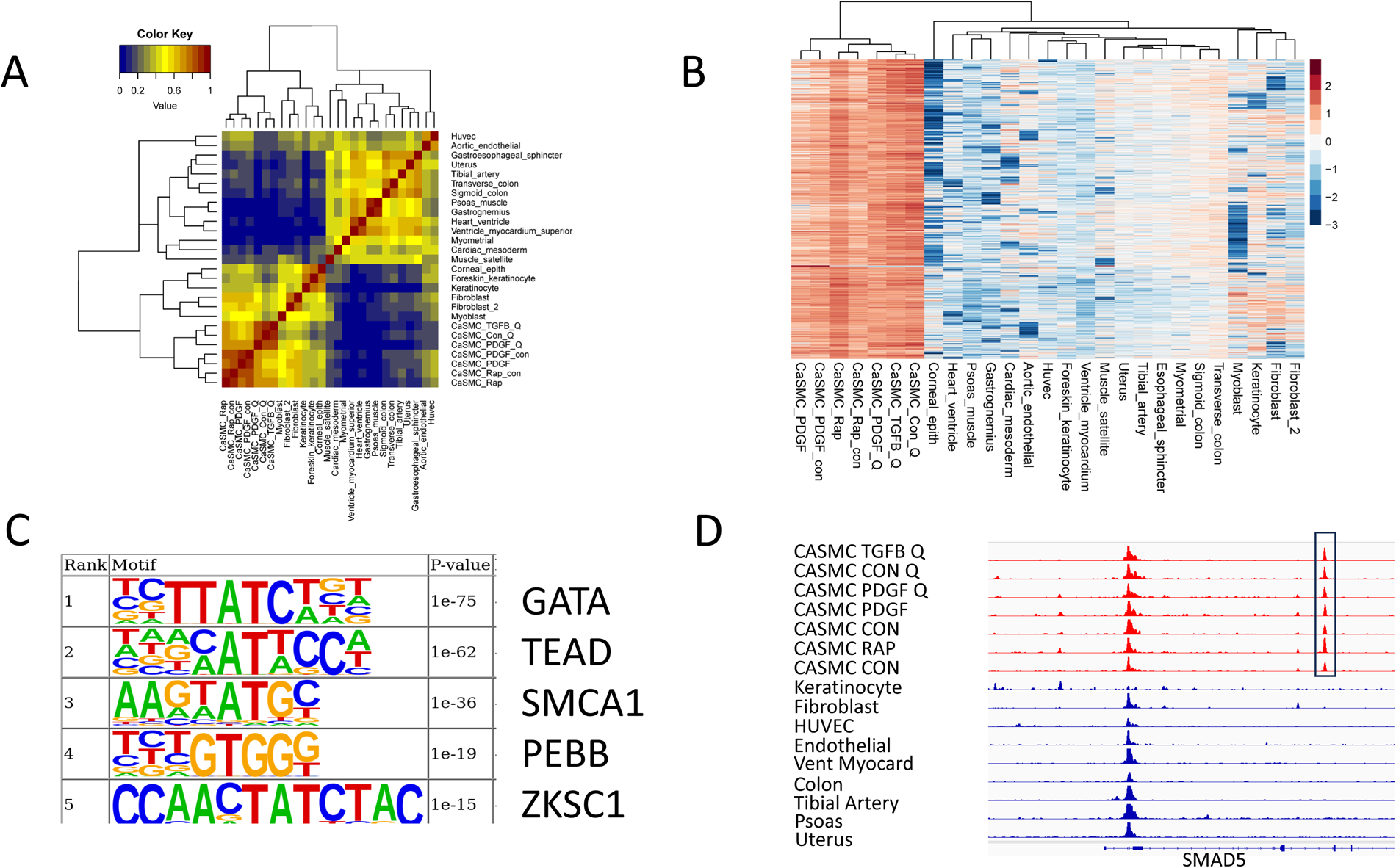
Identification of unique (CASMC)-specific enhancers compared to other cell types. A. Non-promoter ATAC peaks in human cardiac smooth muscle cells compared to other cell types. The heat map shows the Pearson correlation coefficient of read count values. B. Heat map of regions with significantly increased open chromatin in CASMC compared to other cell types; p value cutoff <0.1, 10-fold change. C. Motifs associated with non-promoter ATAC-seq coronary artery smooth muscle cell peaks shown in panel B.

### Enhancer classes and biologically relevant single nucleotide polymorphisms

Genetic variants that modify chromatin accessibility and transcription factor binding are major factors in which genetic variation leads to differences in gene expression ^55–62^. We explored whether SNPs associated with biologically relevant cardiac and vascular traits were enriched in enhancers identified with CASMCs treated with rapamycin or PDGF (Figure 7A). We utilized the set of non-coding SNPs from the GWAS catalog of the NHGRI-EBI (https://www.ebi.ac.uk/gwas/) ^41^ and we collected a data set of cardiac/vascular-associated SNPs. SNP locations were compared to the locations of rapamycin and PDGF enhancers. For active rapamycin enhancers, there was association with 183 SNPs in the GWAS catalog related to cardiac/vascular-associated traits (Supplemental Table S6). For intermediate rapamycin enhancers, there were 129 SNPs related to cardiac/vascular-associated traits. For poised rapamycin enhancers, there were 15 SNPs linked to cardiac/vascular cell traits. Examples of cardiac/vascular-associated SNPs with rapamycin treated CASMC active enhancers are shown in Figure 7B. For active PDGF enhancers, there was an association with 194 SNPs in the GWAS catalog related to cardiac/vascular-associated traits. For intermediate PDGF enhancers, there were 122 SNPs related to cardiac/vascular-associated traits. For poised PDGF enhancers, there were 11 SNPs linked to cardiac/vascular cell traits. For both treatments, active enhancers contained the most cardiac/vascular trait-associated SNPs. The majority of cardiac/vascular trait-associated SNPs were shared between rapamycin- and PDGF-treated enhancers, with some trait-associated enhancers unique to one treatment (Supplemental Table S6). The most common cardiac/vascular trait associated with SMC enhancers was coronary artery disease (Figure 7A).

**Figure 7.**
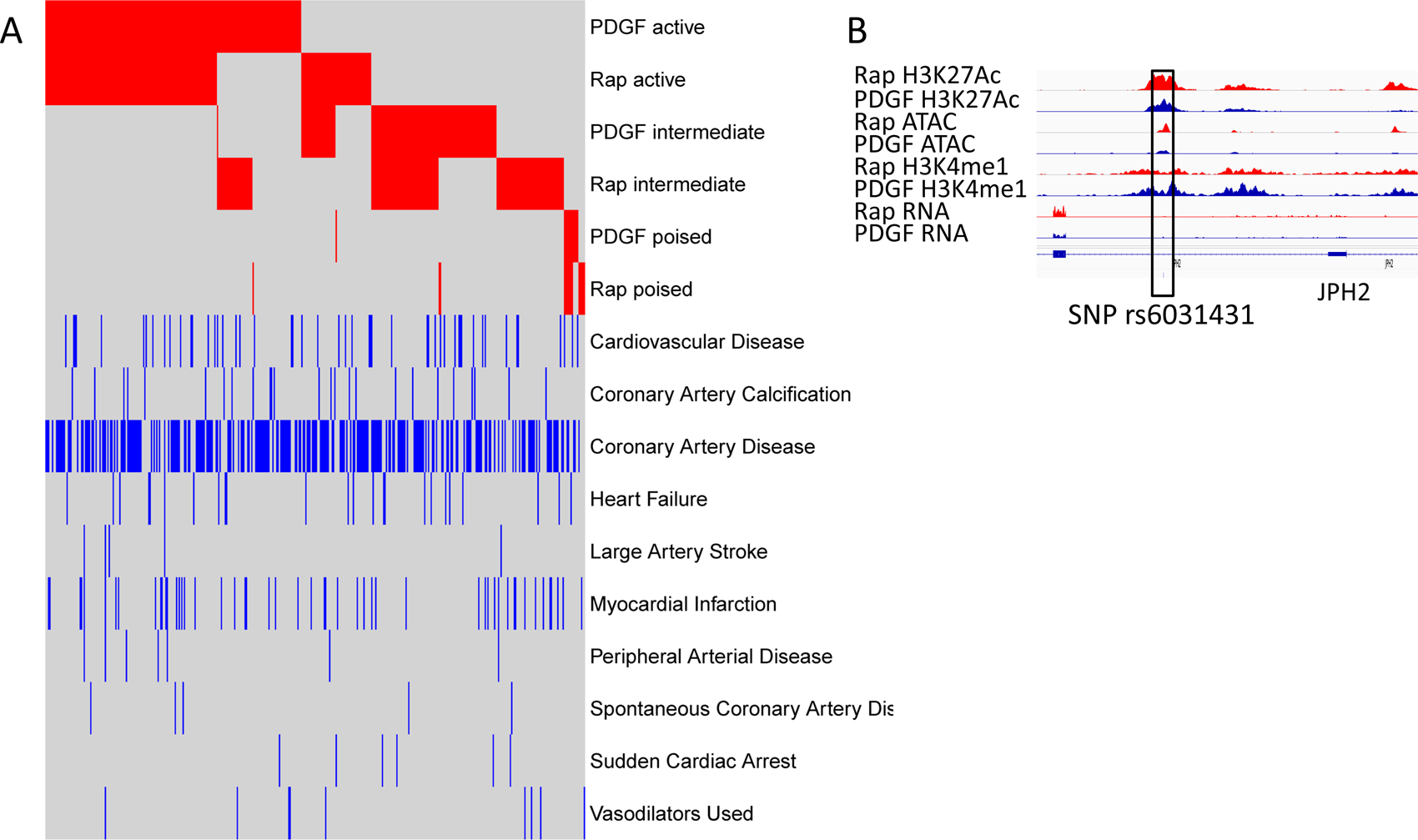
Cardiovascular-associated Genome Wide Association Study Single Nucleotide Polymorphisms (GWAS-SNPs) found in enhancers in rapamycin or PDGF-treated CASMCs. Enhancers in rapamycin- or PDGF-treated cells were mapped onto the GWAS catalog of the National Human Genome Research Institute associated with cardiovascular-associated terms. A. Top ten cardiovascular associated traits are shown with linked enhancer class. The x axis represents the SNPs associated with cardiovascular-associated traits. The top of the heatmap (red) shows each enhancer that contains a GWAS CV-associated SNP by enhancer class. The bottom half of the heatmap (blue) shows the trait associated with each enhancer containing GWAS SNP. Many SNPs were associated with more than one cardiovascular-associated trait. B. Examples of cardiovascular-associated SNPs in enhancers are shown at the JPH2 gene, rs6031431, in rapamycin treated cells and ELL gene in PDGF-treated cells, rs11670056.

### Reporter gene assays of CASMC enhancers

We validated several novel putative enhancer sequences that correspond to SNPs associated with cardiovascular traits by GWAS (Supplemental Table S4). We identified these candidate enhancers as differentially acetylated in CASMC treated with rapamycin, linked to the *CDKN2B* and *JPH2* genes or after treatment with PDGF and linked to the *ELL*, *SLC1A1* and *PDGFD* genes. These sequences were cloned upstream of an SV40 gene promoter-luciferase reporter gene cassette, with the documented *CDKN2B* and *PDGFD* enhancers serving as positive controls ^63^. The resulting plasmids were transfected into CASMCs cells as described. After 48hrs, the cells were harvested, and luciferase activity was analyzed. All five CASMC sequences directed statistically significant (*P* < .05) reporter gene activity between 3 and 12 times over activity of control (Figure 8). Moreover, reporter activity from the *CDKN2B* and *JPH2* sequences was induced by rapamycin treatment of the transfected cells, and reporter activity from the *ELL*, *SLC1A1,* and *PDGFD* sequences were similarly induced by PDGF (Figure 8). These data indicate that these sequences are likely functional enhancers that regulate SMC genes during phenotypic switching.

**Figure 8.**
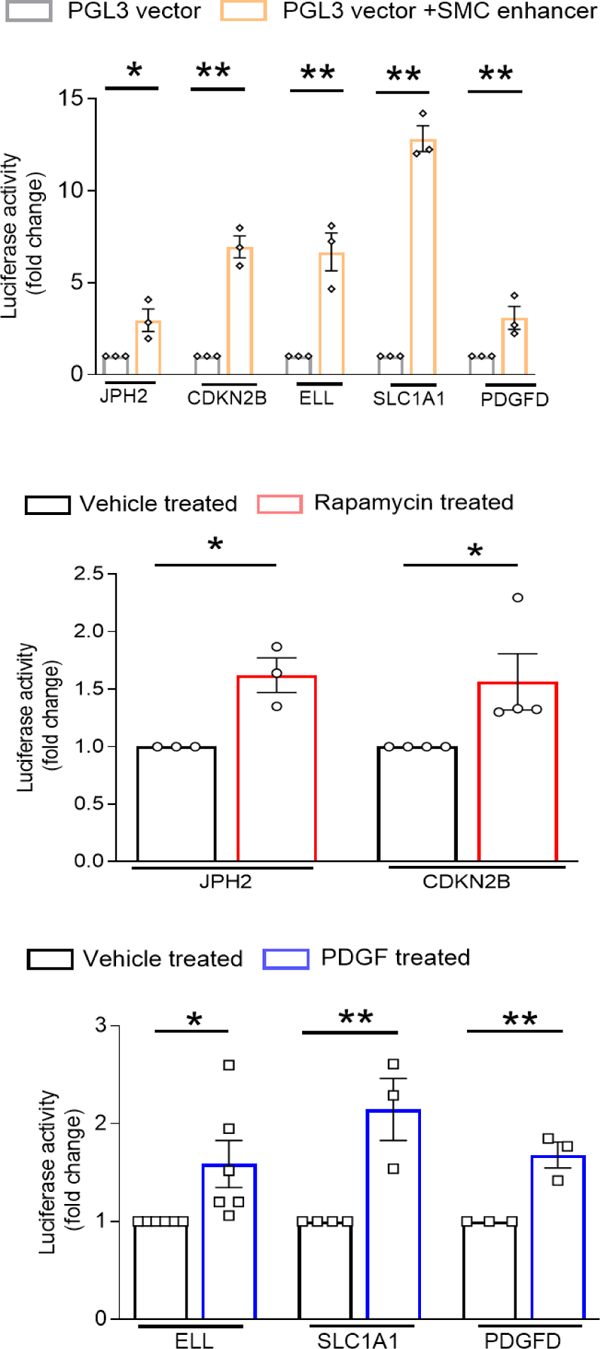
Luciferase reporter gene assays in CASMCs. The activity of novel candidate smooth muscle cell enhancers with known SNPs in luciferase reporter gene assays differentially regulated by rapamycin and PDGF-BB. A) Enhancer activity compared to empty pGL3 vector. Individual reporter gene plasmids were prepared by cloning the indicated SMC enhancer elements upstream of PGL3-SV40 promoter-luciferase reporter gene cassette. These candidate enhancers were identified in our genome wide assays as differentially acetylated by rapamycin in the *CDKN2B* and *JPH2* genes and by PDGF-BB in the *ELL, SLC1A1,* and *PDGFD* genes. The CDKN2B and PDGFD elements served as positive controls. The plasmids were transfected into CASMCs cells as described. After 48hrs, the cells were harvested, and luciferase activity was analyzed. Relative luciferase activity was expressed as that obtained from cells transfected with PGL3-promoter vector with a cloned enhancer compared to that of empty pGL3 vector. B-C) The above plasmids were transfected into CASMCs cells as described. The cells were starved with either 2.5% FBS (for rapamycin) or 0.5%FBS (for PDGF) for 24 hours, then treated with either rapamycin (50 nM) (B) or PDGF (10ng/ml) (C) or vehicle control for additional 48hours. Cells were harvested and luciferase activity was measured. Relative luciferase activity from the Rapamycin- or PDGF-BB-treated reporter plasmid was expressed as relative to that obtained from the same vehicle treated reporter plasmid. The data are the means ± SEM (error bars) of at least 3-4 independent transfection experiments.

A subset of enhancers, called super enhancers or stretch enhancers, important for regulating genes critical for cell-type specific identify, have been described ^19,64^. Super enhancers span large regions of chromatin, have domains of transcription factor binding sites and are marked by significant amounts of H3K4me1 and H3K27Ac modification. In some cell types, disease-associated SNPs are enriched in super enhancers of relevant cell types, suggesting that altered expression of key cell identity genes may contribute to disease phenotype ^64,65^. We identified super enhancers in rapamycin- and PDGF-treated CASMCs by finding regions with the highest levels of clustered, K27 acetylated chromatin (Figure 9) ^19,65^. Super enhancers induced by rapamycin included genes known to be associated with the differentiated SMC phenotype, including *TGFB1*, *TGFB2*, *PTGIS*, and *CALD1.* PDGF-induced super enhancers included *PDGFRA* and *IL1A* (Figure 9), For rapamycin-treated CASMC super enhancers, there was association with 110 SNPs linked to cardiac or vascular traits (Supplemental Table S5) and for PDGF-treated CASMC super enhancers, there was association with 139 SNPs linked to cardiac or vascular traits (Supplemental Table S5). The majority of super enhancer-associated SNPs were associated with the trait “coronary artery disease.” SNPs associated with biologically relevant, disease-associated cardiac or vascular traits were significantly enriched in super enhancers (Fisher’s exact test 5.2e-31) compared to active (1.7e-20) or intermediate (1.4e-10) rapamycin-treated hCASMCs.

**Figure 9.**
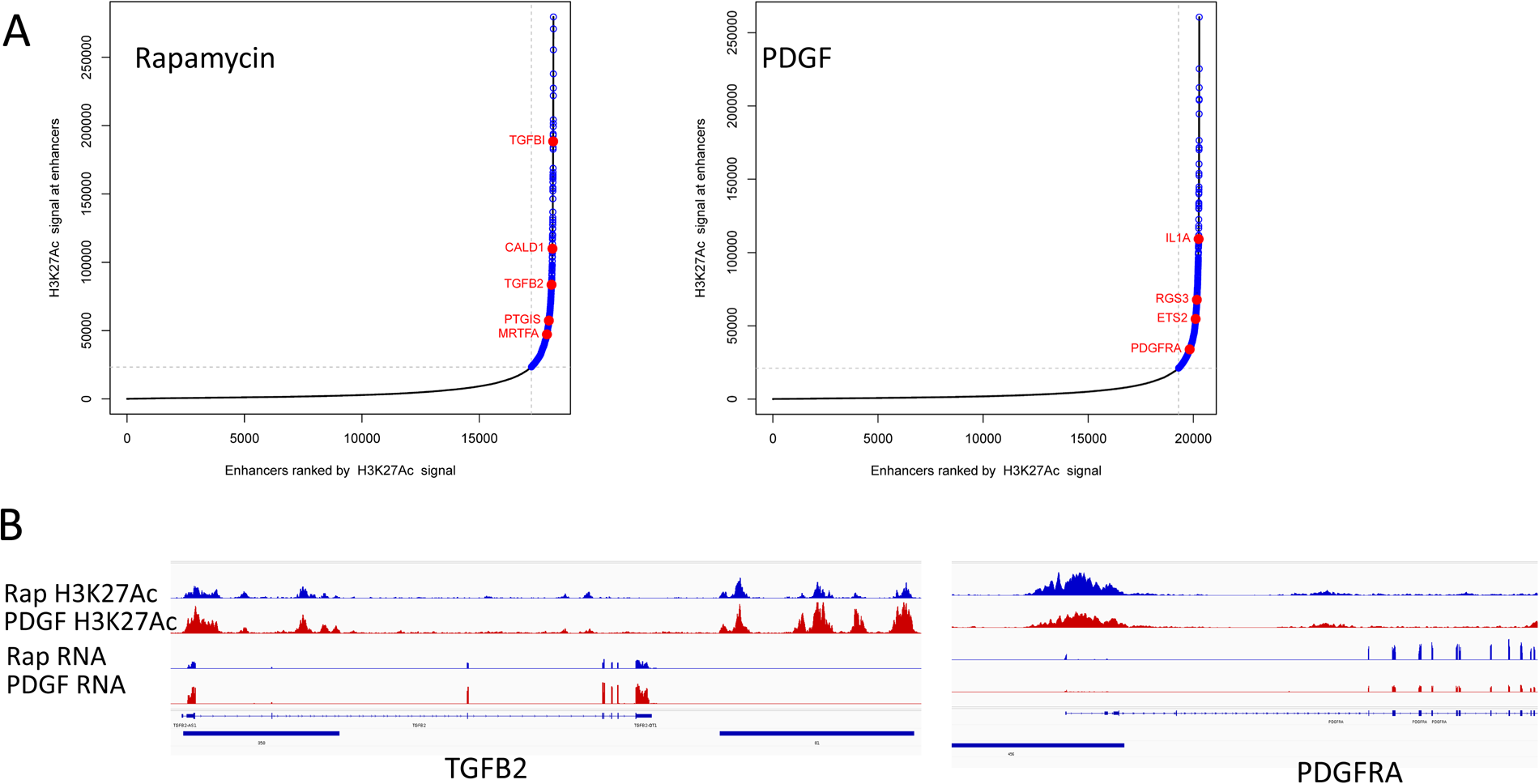
Super enhancers in CASMCs after treatment with rapamycin or PDGF. **A.** Distribution of H3K27Ac normalized ChIP-seq signal across rapamycin or PDGF treated smooth muscle cell enhancers. Select super enhancer-associated genes are labeled in red. **B.** Representative rapamycin-treated smooth muscle cell super enhancer associated with cardiovascular-related SNPs. The super enhancer is denoted by the thick blue line at the bottom of the figure. The associated SNP is shown below the super enhancer line. The track of H3K27 acetylated chromatin is shown above the associated gene locus.

## Discussion

Significant changes in chromatin state dynamics and DNA methylation are typical of cellular development and differentiation ^66–70^. Because differentiated SMC are able to undergo profound state changes, we sought to define differences in the epigenetic landscape in SMC phenotypic switching. Using CASMC treated with rapamycin or PDGF to generate differentiated (contractile) or dedifferentiated (synthetic) phenotypes in culture, we integrated analyses of the transcriptome, accessible chromatin, and ChIP-seq for histone modifications to identify and classify epigenetic states. Our data represent the first detailed epigenetic analyses of differentiated and dedifferentiated CASMCs. Complex, differential patterns of gene expression, promoter state, enhancer location and state, and chromatin configuration were observed after treatment with rapamycin compared to PDGF.

Transcriptomic analyses confirmed that rapamycin and PDGF treatments generated gene expression profiles known to be associated with the contractile and synthetic phenotypes, respectively, and consistent with the metabolic consequences of mTORC1 inhibition, and the mitogenic, pro-migratory effects of PDGF. The ability to pair these changes in gene expression with epigenetic features allows characterization of biological functions, and correlations with large-scale functional datasets such as disease-association studies ^71–73^. These analyses also suggest many exciting new hypotheses that may reveal novel mechanisms underlying SMC phenotypic switching.

We tested the hypothesis that poised promoters, defined as carrying both active (H3K4me3) and repressive (H3K27me3) marks may play an important role in phenotypic switching, with the repressive mark being depleted in response to activating stimuli. Analysis of SMC promoters revealed that only a few hundred promoters changed in state from poised to active in response to rapamycin or PDGF treatment, and these changes did not correlate with changes in gene expression. Instead, a major finding was that there was considerable overlap between the enriched promoter motifs found in the differentiated and dedifferentiated SMC states. This suggests that the well-documented paradigm of SRF binding distinct co-factors at CArG elements in response to extracellular signaling ^10^ may hold true for other transcription factor binding sites in SMC promoters: rather than employing distinctly different types of regulatory elements for alternate fate genes, alterations in signal-responsive transcription factor binding to common sites may allow for rapid change of promoter activity at SMC genes in phenotypic switching.

Other stimuli of CASMC differentiation have been shown to induce epigenetic changes that increase chromatin accessibility and gene transcription, primarily over gene promoters ^63,74,75^. Examples include MYOCD and TGFβ which increase histone 3 acetylation over contractile gene promoters and expression of linked gene expression ^75,76^. In contrast, induction of CASMC de-differentiation by PDGF-BB is associated with chromatin compaction via recruitment of histone deacetylases to contractile gene promoters, decreasing histone acetylation and linked gene expression ^77,78^.

Transcription factor motifs enriched in acetylated regions of promoters of rapamycin treated CASMCs included ZEB, ZBTB7A, and E2F5. Our motif analysis does not discriminate between the binding sites for ZEB1 and ZEB2, but there are data to support a role for either of these transcription factors in SMC phenotypic regulation. ZEB1 (Zinc Finger E-Box Binding Homeobox 1) is a widely expressed member of the zinc finger and homeobox transcription factor family that regulates cell differentiation and a variety of tissue-specific functions, acting as either a transcriptional activator or regulator ^79^. Interestingly, ZEB1 has been implicated in cellular transitions, regulating epithelial-to-mesenchymal transition in cancer cells, and in multiple phenotypic states in macrophages. Recently, decreased levels of Zeb1 in macrophages were shown to increase atherosclerotic plaque formation in mice ^80^. *ZEB1* was linked to coronary artery disease (CAD) in genome wide association studies ^81^, and low ZEB1 expression in human endarterectomies was associated with plaque rupture and cardiovascular events ^80^. Like *ZEB1, ZEB2* has been associated with CAD risk by GWAS ^82,83^ . Studies in mouse atherosclerotic tissues have demonstrated that Zeb2 is transiently expressed in CASMCs during their dedifferentiation and transition to a modulated phenotype, with localization near the fibrous cap. Zeb2 knockdown phenocopied the effects of TGFβ, resulting in similar changes in-gene expression. Zeb2 co-immunoprecipitates with SMAD3 in CASMC, suggesting a potential concerted action of these factors in driving gene expression. Notably, loss of Zeb2 increased SMC contractile gene expression, along with a decrease in synthetic gene expression, including *KLF4*. Combined with our findings, this work suggests that ZEB transcription factors may play important roles in the regulation of coronary artery smooth muscle cell phenotypes ^82^.

ZBTB7A, also known as FBI1 and Pokemon, a transcription factor with C2H2 Krüppel-type zinc fingers acts as a transcriptional repressor of numerous genes involved in cell proliferation and differentiation in many cell types ^79^. Studies of gene expression in human atherosclerotic carotid tissue identified ZBTB7A as a potential regulator of TGFB1 expression in atherosclerosis^84^. E2F5 is a member of the E2F family of transcription factors regulating the coordinated expression of genes involved in cell cycle progression and cell division ^85^. E2F5 is a repressor of gene expression via recruitment of chromatin remodeling factors indirectly to target gene promoters through P130. E2F5 expression is increased in vascular muscle cells in diabetic rats ^86^.

All three of these transcriptional regulators ZEB, ZBTB7A, and E2F5 enriched in the promoters of rapamycin treated CASMCs typically bind in core promoter regions and recruit co-regulatory proteins. These factors have varying functions, with ZEB1 acting as a transcriptional activator, and ZEB1, ZBTB7A, and E2F5 as transcriptional repressors. Interestingly, ZEB and ZBTB7A motifs were enriched in both rapamycin- and PDGF-activated promoters, and the GATA motif was enriched in PDGF induced promoters. There is evidence that GATA-6 may be a bifunctional transcription factor in SMC, driving differentiation ^7^ and dedifferentiation genes ^7,87^ .

These findings demonstrate the complex regulation of programs of gene expression mediated by gene promoters in CASMC phenotype switching.

Our analyses revealed a correlation between H3K27Ac at promoters and gene expression, with differential H3K27 acetylation patterns noted in the contractile versus synthetic states. Our prior work revealed that histone acetyltransferases play key roles in SMC phenotype-specific gene expression. Surprisingly, we found that the highly conserved enzymes p300 and CBP have opposite effects on SMC phenotype, with p300 being inducible by rapamycin and promoting contractile gene expression, and CBP induced by PDGF and promoting synthetic gene expression. Furthermore, these HATs cooperated with other epigenetic regulators. CBP recruited HDACs and deacetylation of contractile gene promoters. In contrast, p300 was found to interact with TET2, and these enzymes function in concert, with each required for the other’s activity at contractile gene promoters ^6^. It will be of interest to extend our studies in the future to determine whether there are distinct p300- and CBP-specific regulatory elements in promoters and enhancers.

The regulation of programs controlling cellular development and differentiation vary temporally, between cell and tissue types, and between species. A major regulator of these are enhancers, elements that recruit regulatory proteins, including transcription factors and chromatin regulators, to influence transcription at distal gene promoters that are frequently associated with phenotypic and disease-associated genetic variants ^41,54,88–91^. Mammalian genomes contain many more enhancers than promoters, and these enhancers play critical roles in controlling gene expression ^53,91,92^. Tissue-specific genes are more dependent on enhancer regulation and exhibit less promoter diversity than housekeeping genes, which are primarily regulated by their promoters with few enhancers in their genomic vicinity ^54^. This observation was true in rapamycin- and PDGF-treated CASMCs where significant changes in enhancer localization, utilization, and state were observed after treatment, and we observed many more differential changes in enhancers than in promoters with changes in SMC phenotype.

Differential H3K27 acetylation analyses identified active enhancers in rapamycin-treated cells with significant enrichment in the transcription factor binding motifs TEAD, SRF, and SMAD. All these factors are known to participate in the pathogenesis of coronary artery disease. SRF and SMADs function distinctly and cooperatively and have been implicated in SMC promoter regulation, so it is perhaps not surprising that they were also found to be major regulators of SMC enhancers. Less is known about TEAD factors, but recent work suggests they also play important roles in SMC. TEAD proteins contain a YAP-binding domain and a DNA binding domain but require a coactivator for its transcriptional activity ^93^. Variation in coactivator identity determines patterns of gene expression. Major TEAD partners are YAP or TAZ, key regulators of the Hippo signaling pathway. This pathway has been implicated in inflammation associated with cardiovascular diseases such as myocardial infarction, cardiomyopathy, and atherosclerosis ^94^. The TEAD/YAP axis has been shown to regulate gene expression in cardiomyocytes, macrophages, and myocardial fibroblasts. TEAD1 has been shown to promote mTORC1 activity and a synthetic phenotype in SMC. Accordingly, smooth muscle-specific knockdown of TEAD1 prevents intimal hyperplasia in a mouse model through inhibition of PDGFRβ expression and VSMC proliferation^95^. Our data indicate a role for this axis in remodeling of the CASMC epigenome after rapamycin treatment.

Genetic variants that modify chromatin accessibility and transcription factor binding are major factors in which genetic variation leads to differences in gene expression ^56–58,62^. These genetic variants are enriched in functional epigenomic elements, frequently in gene enhancers, influencing quantitative traits and disease phenotypes. SNPs linked to cardiovascular disease phenotypes by genome wide association studies were highly enriched in enhancers from both rapamycin- and PDGF-treated cells, and these enhancers were located near genes highly expressed in CASMCs or involved in cardiovascular function. Biologically relevant SNPs were most enriched in active enhancers and super enhancers, with many mapping to sites previously associated with cardiovascular disease in genome-wide association studies. However, this study identified numerous biologically relevant SNPs not previously linked to known functional regulatory elements in CASMCs. These data provide a significant resource for studies of cardiovascular disease and the role of CASMC in normal and perturbed states.

Several groups have performed integrative epigenetic analyses in CASMC to identify genes linked to coronary artery disease (CAD) ^63,90,96–100^, but these studies have not performed a comprehensive mapping of promoters and enhancers in distinct phenotypic states. One study used brief six-hour TGFβ or PDGF treatments combined ATAC-seq with GWAS data to identify regulatory regions associated with CAD risk ^63^. Another study applied H3K27ac HiChIP to generate high-resolution contact maps of active enhancers and target genes, with a primary focus on T cell differentiation. This study included an analysis of CASMC in a static condition, without assessing changes with phenotypic switching ^101^. Our study is the first to compare CASMC in a fully differentiated or dedifferentiated state to identify state-specific epigenetic regulatory signatures and motifs. This information will be important as CAMSC are a major target in CAD and coronary artery revascularization. SMC arise from different developmental origins in distinct regions of the vasculature ^102^. We suspect that many of our findings may apply in other vessels, but it is possible that some patterns may be specific to CASMC.

SMC can undergo multiple fate transitions, including toward fibroblast, macrophage-like, chondrocyte-like, and even adipocyte-like phenotypes ^82^. It will be of interest to compare epigenetic mechanisms that control these various transitions. Unbiased comparative integrated analyses may reveal novel motifs and transcription factors that influence these distinct states which may provide new avenues for prevention and intervention in cardiovascular disease.

In summary, this study is the first to identify and characterize enhancers and super enhancers regulated by rapamycin and PDGF in human CASMCs. Integrative analyses identified complex patterns of promoter and enhancer state in differentiated and dedifferentiated CASMCs. Single nucleotide polymorphisms linked to cardiovascular disease phenotypes by genome wide association studies were highly enriched in enhancers, particularly super enhancers, from both rapamycin- and PDGF-treated cells, and these enhancers were located near genes highly expressed in CASMCs or involved in cardiovascular function. These data provide a significant resource for epigenetic studies of CASMC in normal and perturbed states as well as for genetic studies of cardiovascular disease.

## Nonstandard Abbreviations and Acronyms

hCASMC: Human coronary artery smooth muscle cells
ChIP-seq: Chromatin immunoprecipitation and sequencing
TSS: Transcriptional start site
TGF-β: Transforming growth factor
mTORC1: Mammalian target of rapamycin complex 1
PDGF: Platelet derived growth factor
SNP: Single nucleotide polymorphism

