## Supplemental Table 1 for "Epigenetic Remodeling in Human Coronary Artery Smooth Muscle Cell Phenotypic Switching"

Supplemental Table S1. Primers for Amplification of Candidate Enhancers

*JPH2*
Forward
5’-AGCA GGTACC AAATGAAAGGGAAAAGTGAACAGGCAG-3’
Reverse
5’-AGCA CTCGAG CAGTGAGGAGGCAGTCCAAAG-3’

*CDKN2B*
Forward
5’- AGCA GGTACCGGGAAAATTGAGGCAGGTC-3’
Reverse
5’- AGCA CTCGAGCAGGGGGTTGGCTGTCTAC-3’

*ELL*
Forward
5’- AGCA GGTACC GACAGCAATGCCTTTGGG-3’
Reverse
5’- AGCA CTCGAGCTTCTCAGCCCCTGCAC-3’

*SLC1A1*
Forward
5’- AGCA GGTACC GCTACGCAAGCTGAGGTTGTTCAC-3’
Reverse
5’- AGCA CTCGAGCACTCCAAAGGAAGCAGTTGCC-3’

*PDGFD*
Forward
5’- AGCA GGTACC GCTGGACTATGTAGAATTCATCCAGATGGG-3’
Reverse
5’- AGCA CTCGAG GGGAGTTTCTATTGTTCTGCCCAGG-3’
